# CC^+^: A Searchable Database of Validated Coiled coils in PDB Structures and AlphaFold2 Models

**DOI:** 10.1101/2023.06.16.541900

**Authors:** Prasun Kumar, Rokas Petrenas, William M. Dawson, Hugo Schweke, Emmanuel D. Levy, Derek N. Woolfson

## Abstract

α-Helical coiled coils are common tertiary and quaternary elements of protein structure. In coiled coils, two or more α helices wrapped around each other to form bundles. This apparently simple structural motif can generate many architectures and topologies. Understanding the variety of and limits on coiled-coil assemblies and their sequence-to-structure relationships impacts on protein structure, design, and engineering. Coiled coil-forming sequences can be predicted from heptad repeats of hydrophobic and polar residues, ***hpphppp***, although this is not always reliable. Alternatively, coiled-coil structures can be identified using the program SOCKET, which finds knobs-into-holes (KIH) packing between side chains of neighboring helices. SOCKET also classifies coiled-coil architecture and topology, thus allowing sequence-to-structure relationships to be garnered. In 2009, we used SOCKET to create a relational database of coiled-coil structures, CC^+^, from the RCSB Protein Data Bank (PDB). Here we report an update of CC^+^ following the recent explosion of structural data and the success of AlphaFold2 in predicting protein structures from genome sequences. With the most-stringent SOCKET parameters, CC^+^ contains ≈12,000 coiled-coil assemblies from experimentally determined structures, and ≈120,000 potential coiled-coil structures within single-chain models predicted by AlphaFold2 across 48 proteomes. CC^+^ allows these and other less-stringently defined coiled coils to be searched at various levels of structure, sequence, and side-chain interactions. The identified coiled coils can be viewed directly from CC^+^ using the Socket2 application, and their associated data can be downloaded for further analyses. CC^+^ is available freely at http://coiledcoils.chm.bris.ac.uk/CCPlus/Home.html. It will be regularly updated automatically.

**FOR THE BROADER AUDIENCE:** Protein assemblies and protein-protein interactions are key to all biological processes. α-Helical coiled coils are one of the most common modes of directing and stabilising these interfaces. Here, we report an updated CC^+^ database of structurally validated coiled coils from experimental protein structures and AlphaFold2 models. CC^+^ contains many thousands of coiled-coil structures and models, associated parameters, and sequences. It enables the compilation of rich datasets for advancing protein structure, design, and engineering research.

## 1 INTRODUCTION

Protein-protein interactions (PPIs) are critical for all biological processes.^1^ There are various classes of PPI involving the common protein secondary structure elements—α helices and β strands—less-well-defined turns, loops, and intrinsically disordered regions, and many types of protein tertiary structure. Gathering sequence and structural data on PPIs^2-5^ is important and necessary if we are to understand protein networks, target them for biomedical applications, and exploit them in synthetic biology and biotechnology.

One class of PPI that is well defined both at the sequence and structural levels is the α-helical coiled coil (CC). As a result, CCs can be readily identified and examined. In turn, this provides insight into protein structure and function, and a solid basis for protein design and engineering. Indeed, structural, design, and engineering studies of CCs are relatively mature and have been reviewed extensively.^6-12^ Therefore, this introduction focusses on the most-relevant points that underpin the work presented herein. One thing lagging behind these advances is an understanding of the biological diversity and functions of the many natural CCs.^12^ We hope that the revised database of CC structures and associated data presented here will help reach a complete understanding of this particularly widespread and diverse class of PPI.

CCs are assemblies of two or more α helices that wrap around each other to form rope-like, or supercoiled, helical bundles. At the most basic level, these assemblies are encoded by 7-residue (heptad) sequence repeats in which hydrophobic (***h***) residues are spaced alternately 3 and 4 residues apart. The intervening residues are often polar (***p***) resulting in ***hpphppp*** repeats usually labelled ***abcdefg***. Combined with the 3.6 residues per turn of the α helix, these patterns encode amphipathic helices with distinct hydrophobic (***a*** + ***d***) and polar faces. In aqueous media, multiple copies of such helices come together to bury their hydrophobic faces and form hydrophobic cores that stabilises the assemblies. The helices supercoil around each other because the average hydrophobic spacing of 3.5 residues falls short of the fixed 3.6 residues per turn of the α helix. Moreover, and because of this combination of sequence repeat and α-helical geometry, residues at ***a*** and ***d*** form *knobs* that can insert into *holes* formed by four residues of a neighbouring helix (*e*.*g*., ***d-g-a-d*** or ***a-d-e-a***, respectively, in parallel CCs). This so-called knobs-into-holes (KIH) packing is the true signature of CC structures and assemblies.

However, this apparent simplicity masks underlying complexities.^7,13^ For instance, although heptad sequence repeats are the most common, others based on 3,4-spacings of hydrophobic residues are possible, and these lead to different CC supercoils. Also, as well as different CC oligomers from dimers upwards, the component helices of CC assemblies can be in all-parallel, antiparallel, or mixed arrangements. And, though many CCs are homo-oligomers, hetero-oligomers are also common. Finally, as well as these oligomeric CCs, *i*.*e*. quaternary structures or PPIs, many CCs are formed within single chains, *i*.*e*. they form parts of tertiary structures. Access to complete and robust databases of CCs would help assess the breadth of CC structural space, and the interrogation of CCs *en masse* would give a deeper understanding of sequence-to-structure/function relationships. In turn, this would aid structural molecular biology, chemical and synthetic biology, protein design, and other fields and applications.

Some time ago, we wrote the program SOCKET to identify KIH packing between α helices in 3D coordinates of protein structures, *i*.*e*., RCSB PDB files.^14,15^ SOCKET has allowed us to identify CCs in the whole of the PDB and create a relational database of CC structures, CC^+^.^16^ Using these resources, we have been able to classify CCs in a Periodic Table of Coiled Coils^17^ and a graph-based Atlas of Coiled Coils ^18^. Others have used SOCKET to generate similar databases and useful resources for collating and analysing CC structures; notably, SamCC-Turbo.^19^ Web-based resources for predicting, analysing, and categorising CCs more widely have been reviewed elsewhere.^12^

Recently, we have updated SOCKET to Socket2.^20^ This includes improvements in the SOCKET algorithm itself to capture CC structures that SOCKET could not find, and a new visualiser to allow users to view and analyse identified CCs directly in real time. Here we describe an update and overhaul of the CC^+^ database using Socket2. Moreover, we have taken this opportunity to include CCs identified by Socket2 in both the experimentally validated structures of the RCSB PDB and those found in tertiary structures predicted by AlphaFold2^21^ collated in EMBL-EBI AlphaFold Protein Structure Database from 48 genomes.^22^ We envision that the new CC^+^ database and its associated tools will be of use to expert and non-expert users interested in all aspects of CC biology, structure, and design.

## 2 DESIGN, ARCHITECTURE, AND POPULATION OF THE CC^+^DATABASE

The new CC^+^ Database has three components: First, an automatically updated backend houses the core data on the identified CC structures/models and their associated sequences and structural parameters. Second, as described in the next section, an accessible frontend allows users to access these data using a wide range of search parameters and criteria. Third, following user-defined searches, CC^+^ can generate search-specific data on the fly, including: position-specific scoring matrices (PSSMs) from the selected sequences; and models of the CC regions in context of the whole protein structure, which can be visualised and interrogated using the interactive GUI of Socket2.^20^

The design and architecture of the backend borrow from the original database reported in 2009:^16^ *i*.*e*., it is an updated rather than a completely rewritten resource. The process of populating this is outlined in Figure 1 and described below.

**FIGURE 1.**
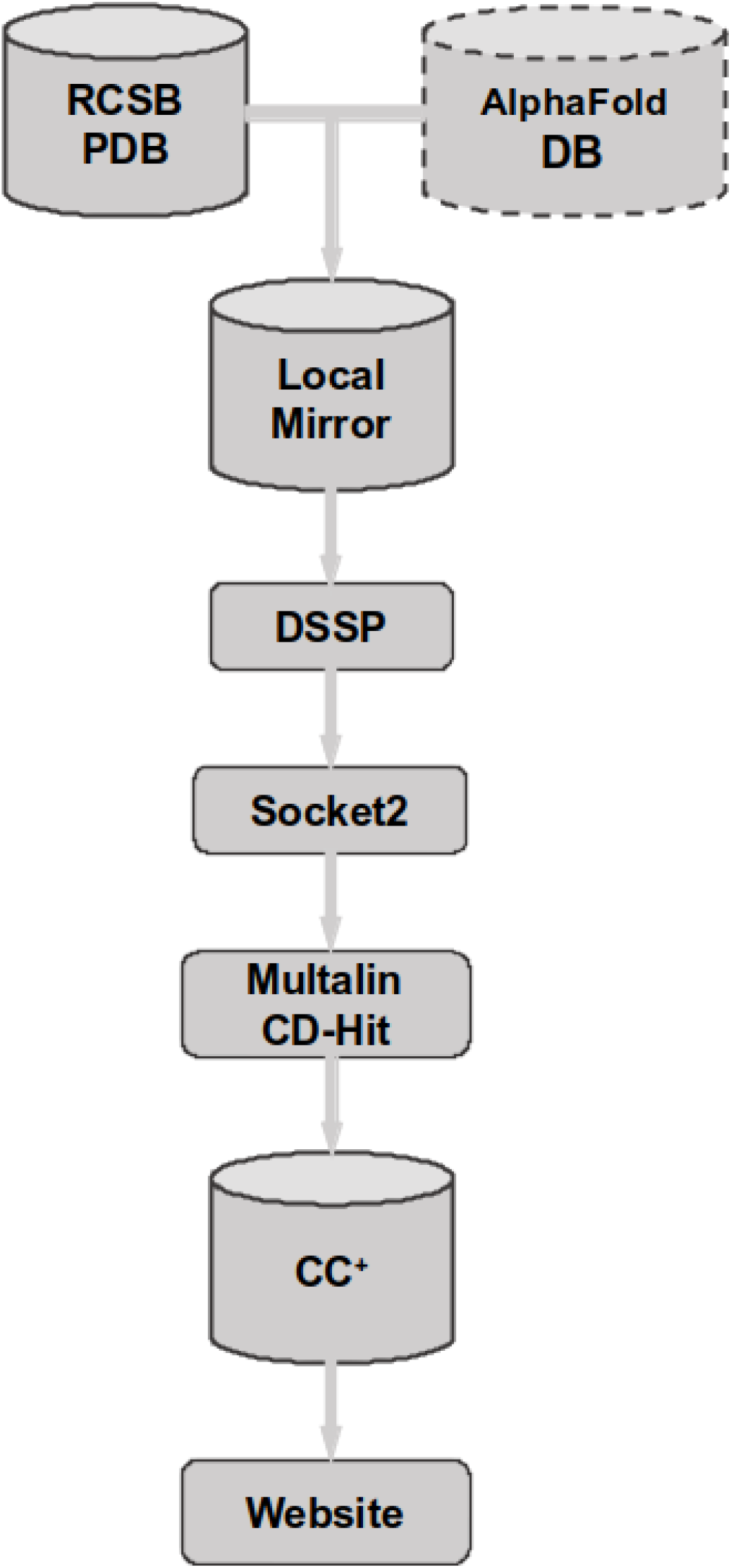
Flow chart detailing the process of compiling the CC^+^Database and website. The CC^+^database is compiled from the structures from RCSB PDB^14^and models from the AlphaFold Protein Structure Database.^22^These files are scanned using DSSP^23,24^and then Socket2^20^to identify KIH interactions and assign CCs. The organized output is stored in a MySQL database comprising tables of CCs and associated data. MultAlin^25^and CD-HIT^26,27^are used to align sequences and calculate redundancy, respectively. A user-friendly website has been developed to provide easy access to and searching of the stored data at http://coiledcoils.chm.bris.ac.uk/CCPlus/Home.html.

PDB-formatted files from the January 2023 release of RCSB PDB (PDB)^14^ and the AlphaFold Protein Structure Database of predicted models for 48 genomes^22^ were downloaded. For the PDB structures, the corresponding asymmetric and biological units were also considered in the following process. DSSP^23,24^ and Socket2^20^ were used to identify α-helical regions and to assign CCs, respectively. Multalin^25^ was then used to determine if partnering helices had the same or different sequences. CD-Hit^26,27^ was used to categorize sequences into four groups based on redundancy: ≤50% identity, ≤70% identity, Non-redundant, and Redundant. As described below, these categories can be used to focus user-defined searches. Biopython^28^ was employed to generate PDB files for just the CC regions. CC assignments and associated data were stored in a MySQL database for easy access and searchability. The full set of these assignments, or ‘default structures’, are summarized in Figure 2, where they are broken down according to the number and orientation of α helices in each CC.

**FIGURE 2.**
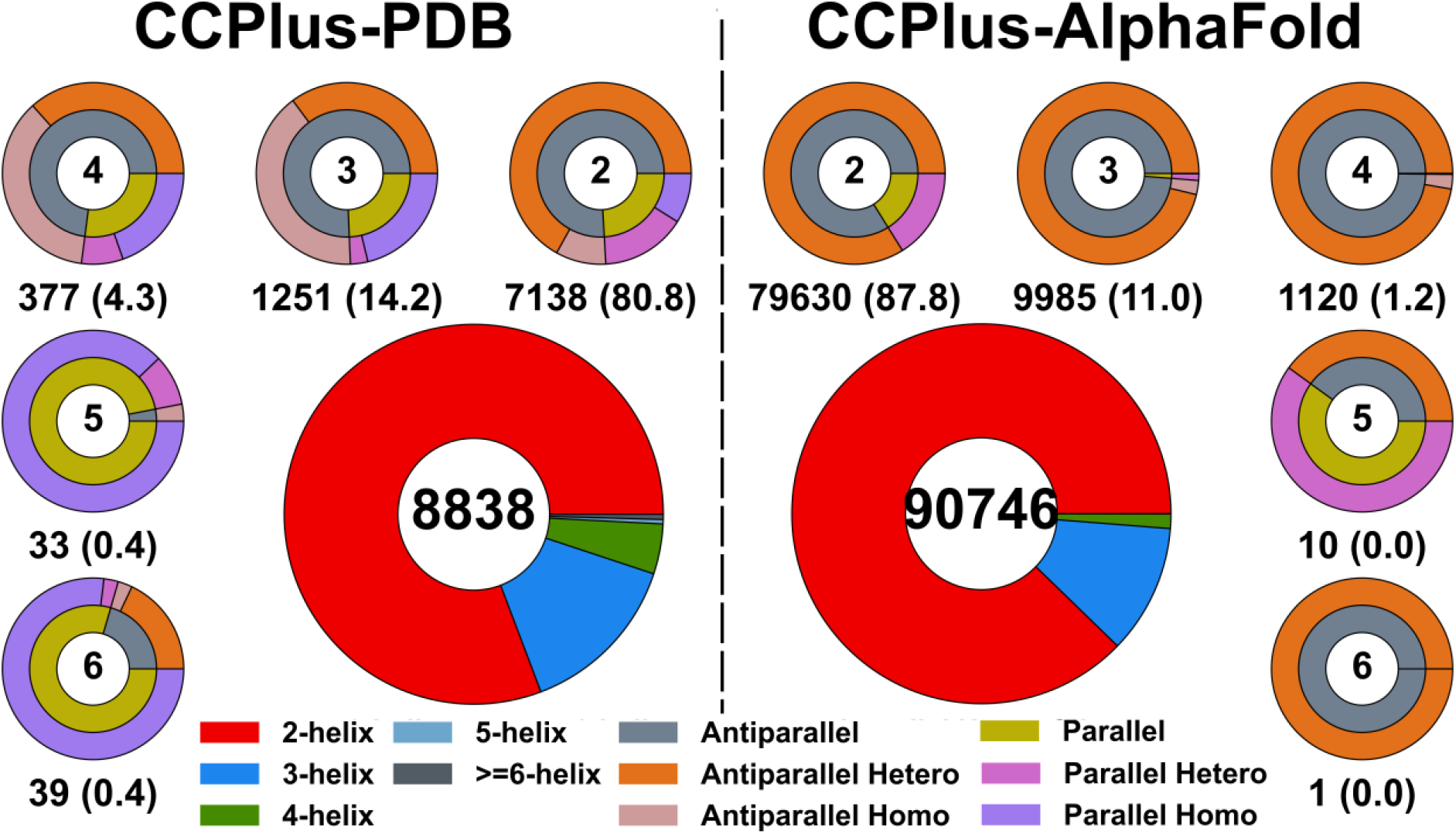
A summary of the CC composition of the CC^+^Database as of January 2023. The CCs are derived from either protein structures deposited in RCSB PDB or predicted by AlphaFold2 and hosted at the EBI. The two large doughnut charts give the total numbers of CCs found in the two sub-databases using default parameters of 7 Å SOCKET packing cut-off and 70% sequence identity. Here the coloring is by number of α helices in the CC assembly; *i*.*e*., 2, 3, 4, 5 and ≥6 helices. The smaller peripheral doughnuts give the breakdown of these CCs based on the number of helices (centered numbers). Here the coloring is by topology; *i*.*e*., antiparallel, parallel, etc. The numbers below each doughnut indicate the total number of CCs and percentages (bracketed) for each class. Note: there are relatively few CCs with six or more α helices, which make up less than 1% of the returned structures.

The user-accessible frontend of CC^+^, was developed using HTML, JavaScript, and CSS, and the backend using CGI/Perl, HTML, JavaScript, and CSS. The MySQL database was used to create the associated sequence and parameter tables. For on-the-fly sequence analyses, Python modules were used to generate PSSMs and perform χ^2^ tests. Matplotlib^29^ and PyMol^30^ were used to generate plots and figures for protein structures, respectively. The CC^+^ Database is available at http://coiledcoils.chm.bris.ac.uk/CCPlus/Home.html as part of a Linux, Apache, MySQL, CGI, JavaScript, and HTML stack.

## 3 USING THE CC^+^DATABASE

### 3. Overview

The CC^+^ Homepage provides links to two form-based ‘Dynamic Interface’ tabs, one each for customizable searches of coiled coils in PDB structures and AlphaFold2 models, CCPlus-PDB (Figure 3) and CCPlus-AlphaFold, respectively. These are described in detail below. The ‘Statistics’ tab provides users with quick-glance summaries of the content of the current version of the database in terms of the number of CCs categorised as intra-(same chain) and inter-molecular (many chains), the number of helices/oligomeric state in the CC assemblies, and the arrangement of helices within these (parallel or antiparallel). These summaries can be tailored by users selecting the Socket2 cut-off value (7, 7.5, 8, 8.5 or 9 Å) and the sequence redundancy or similarity (50%, 70%, Non-identical, Redundant) used to find the CCs. Finally, the ‘Documentation’ tab provides users with comprehensive descriptions of Socket2^20^, using the Dynamic Interface, where searches can go wrong, and how data can be downloaded and used.

**FIGURE 3.**
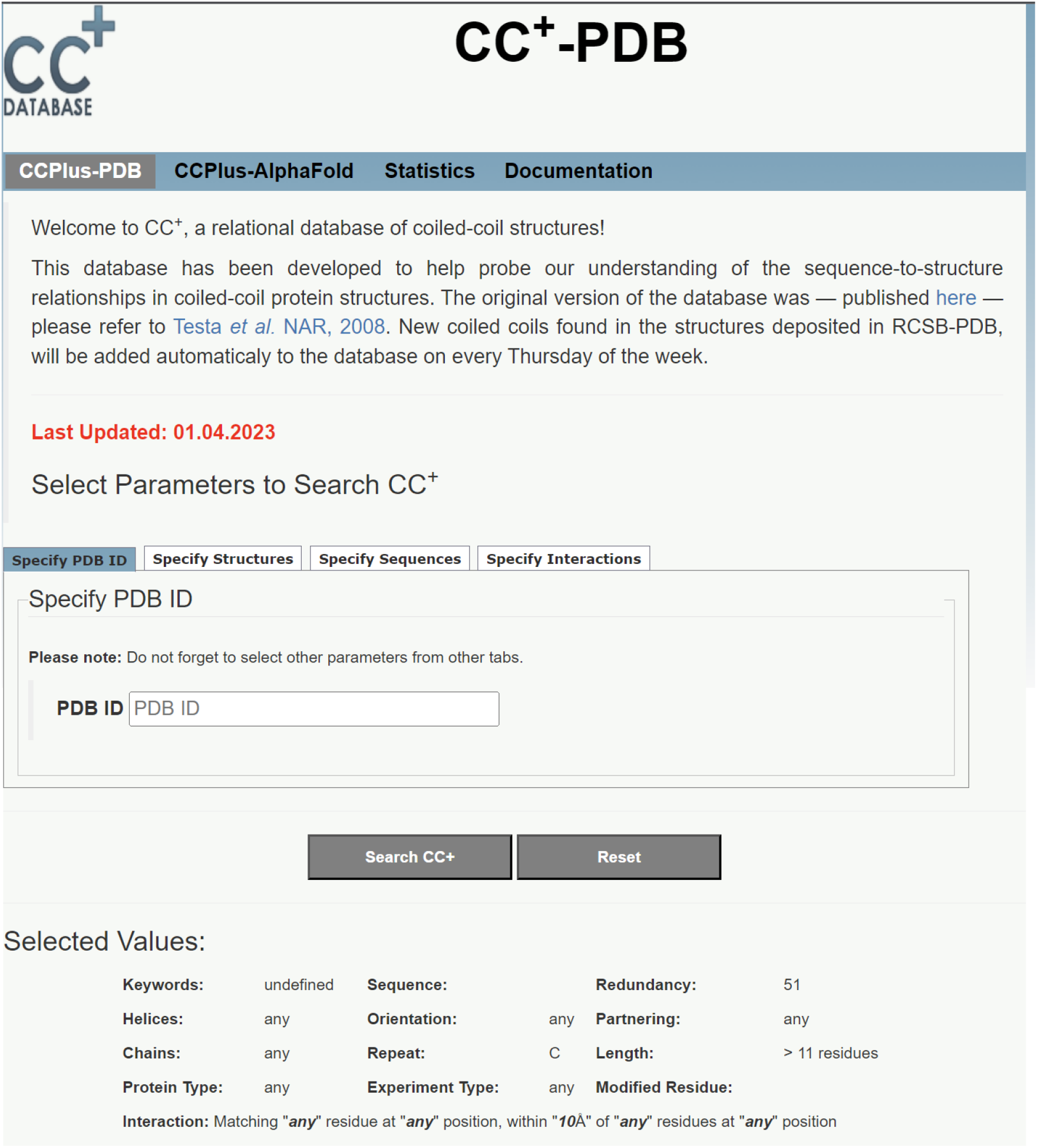
Entry page for the CCPlus-PDB ‘Dynamic Interface’. The page contains a brief description and last date of the update. The page also provides four tabs containing different parameters to explore the database. These options are discussed below. The search can be initiated by clicking Search CC^+^ button, while `Reset` button resets the parameters to their default values. The last section gives the current selected values for different parameters.

### 3. Searching the Dynamic Interfaces

A major advance in protein science since the launch of the CC^+^ database in 2009 has been the success of AlphaFold2 in predicting protein structures^21^ and its application to 48 complete genomes.^22^ Therefore, along with CC structures found by Socket2^20^ in the RCSB PDB,^31^ for the updated CC^+^ we have included those found in AlphaFold2 models predicted from these 48 proteomes.^22^ However, to avoid confusion between experimental structures and predicted models, and to give users control over how they use (separate or combine) these data, we have separated the backend data and frontend searches of the two groups *via* the CCPlus-PDB and CCPlus-AlphaFold ‘Dynamic Interface’ tabs. As detailed below, some of the subtabs used to search these are common to both parts of the CC^+^ Database, but others are unique to each arm of the database. In all cases, searches are initiated by clicking the Search CC^+^ button, and default values for each of the parameters can be regained with the `Reset` button. Searches using default values include any number of canonical helices that are over 11 residues in length, in any orientation, with any type of partnering, from any number of chains, and with 50% sequence identity or less.

#### Specify PDB IDs

This subtab only applies to CCPlus-PDB. It can be used to find any CCs in a chosen PDB file. This can be used in conjunction with other search parameters. However, to avoid missing any CCs, we advise setting the `Redundancy` parameter to ‘redundant’ and leaving the other subtabs at their default settings in the ‘Specify Structures’ subtab.

#### Specify Structures

This subtab has undergone major updates. For both the CCPlus-PDB (Figure 4A) and CCPlus-AlphaFold (Figure 4B) tabs, a slider allows users to choose the Socket2 cut-off value for identifying CCs. The default value is 7 Å, which we recommend using, as the other values (7.5, 8, 8.5, and 9 Å) are increasingly less stringent and may pull in non-coiled-coil regions.

**FIGURE 4:**
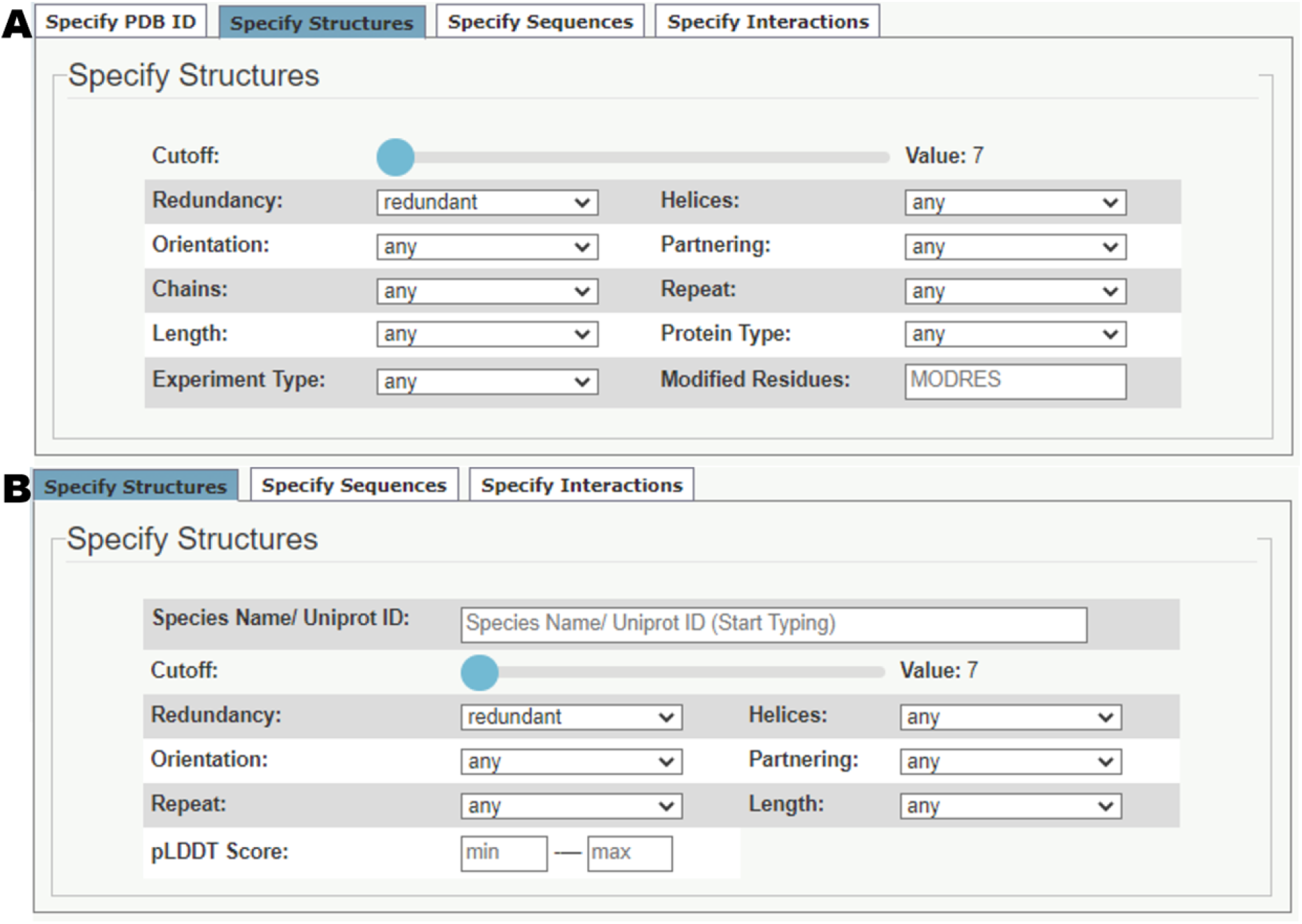
‘Specify Structures’ subtab from (A) CCPlus-PDB and (B) CCPlus-AlphaFold ‘Dynamic Interface’ tabs. The default values for each parameter are shown.

The ‘CCPlus-PDB’ tab (Figure 4A) offers several search parameters for CCs, including: `Redundancy`, as introduced above; the number of α `Helices` and their relative `Orientation`; whether the `Partnering` helices have the same (homo-mers) or different (hetero-mers) sequences; whether the helices are from the same or different polypeptide `Chains`; if the underlying CC sequence `Repeats` are heptad or non-heptad based; and the minimum `Length` of the CC helices. In addition to these options adopted from the original CC^+^, users can now specify the `Protein Type`(membrane or globular); the `Experiment Type` used to solve the parent structure; and requesting specific `Modified Residues`, *i*.*e*., non-proteinogenic residues. When `Experiment Type` is defined a `Resolution` range can be added by the user.

As AlphaFold2 predicted models are for single chains only and contain only the 20 standard proteogenic residues, the ‘CCPlus-AlphaFold’ tab (Figure 4B) offers similar search parameters to the above but without options for the number of `Chains`, `Protein Type`, `Experiment Type`, and `Modified Residues`. These searches can also be filtered by a min—max `pLDDT Score` for the predicted CC regions.

#### Specify Sequences

Searches can be defined further using this subtab (Figure 5A) to find CCs that contain specified sequences or sequence patterns. Users can enter plain text for a query sequence or use PROSITE^32^ notation to find sequence patterns. Also, these sequences and patterns can be requested to fall at specified positions of the heptad repeats using the ***a – g*** heptad notation.

**FIGURE 5:**
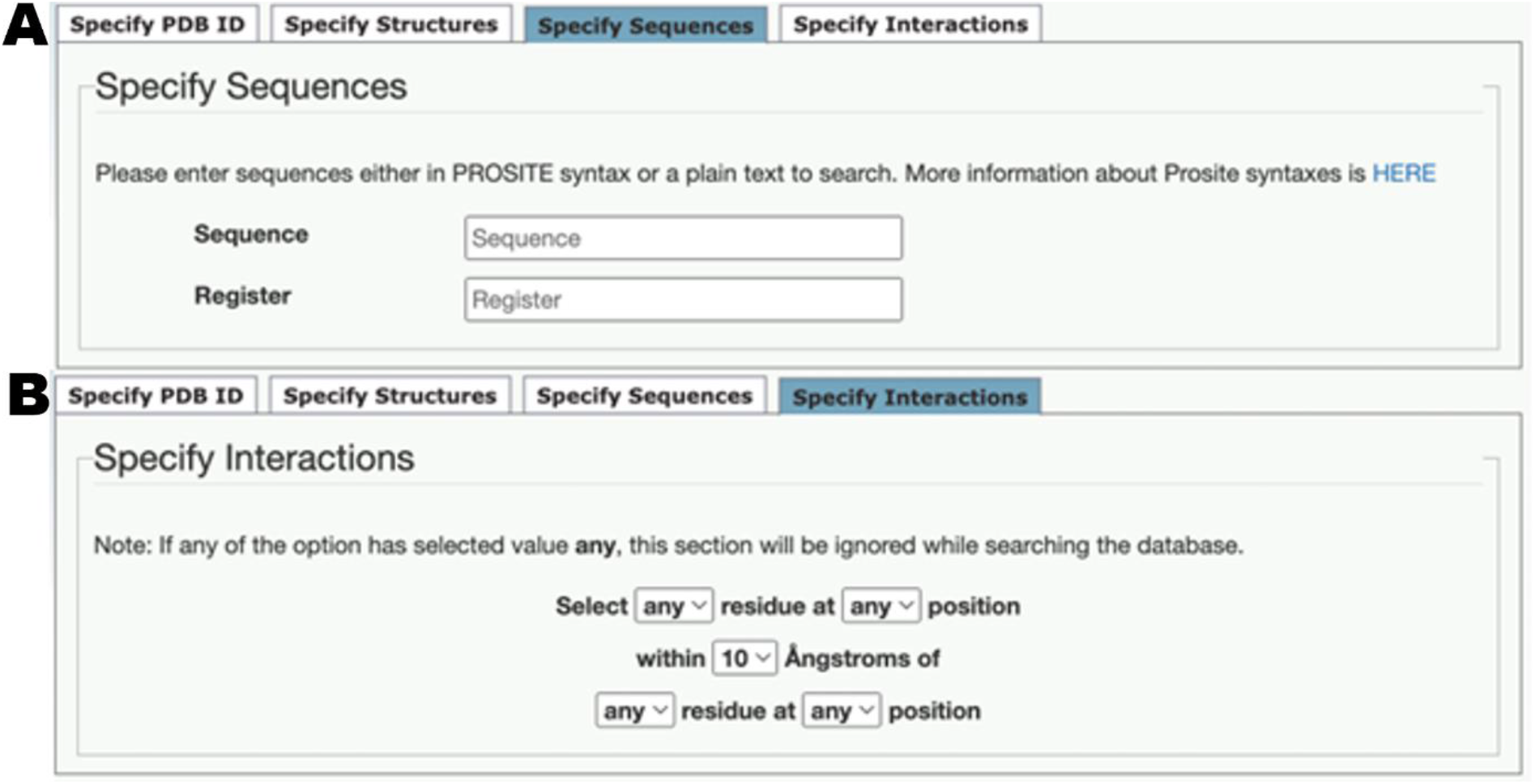
(A) ‘Specify Sequences’ and (B) ‘Specify Interactions’ subtabs for CCPlus-PDB and CCPlus-AlphaFold. Default value of each parameter is also shown.

Note: The `Register` field of this subtab can be used to search for non-canonical repeats. For instance, 11-residue, hendecad repeats can be found by entering “***abcdefgdefg***”. This is because Socket2 locates KIH interactions and *then* assigns them as ***a – g*** sites only, as heptad repeats predominate in CC sequences and structures. In the case of hendecad repeats, the ***hijk*** positions are analogous to an addition of ***defg*** to a heptad repeat.

#### Specify Interactions

Finally, searches can be defined even further using this subtab (Figure 5B), which allows users to identify residue-residue interactions within a specified distance in a CC search. This has the option to specify the register positions of the interacting residues. Thus, this feature allows the compilation of CC datasets with potential residue-residue interactions that underpin sequence-to-structure relationships in CCs.

### 3.3 Displaying and using results from CC^+^searches

On completion of a search, the results are displayed in a new page, Figure 6. Here users can choose several options to display and analyse the data. Each ‘Results page’ has four main features: (i) ‘Selected Values for the Current Search’, which simply tabulates the current selection criteria and their values; (ii) the ‘Results’ section itself, which is explained below; (iii) an associated ‘Gallery’ of cartoons for the CCs returned, which allows quick access to their structures and sequences; and (iv) a ‘Search Again’ tab, which allows the user to search the database with a new set of parameters.

**Figure 6:**
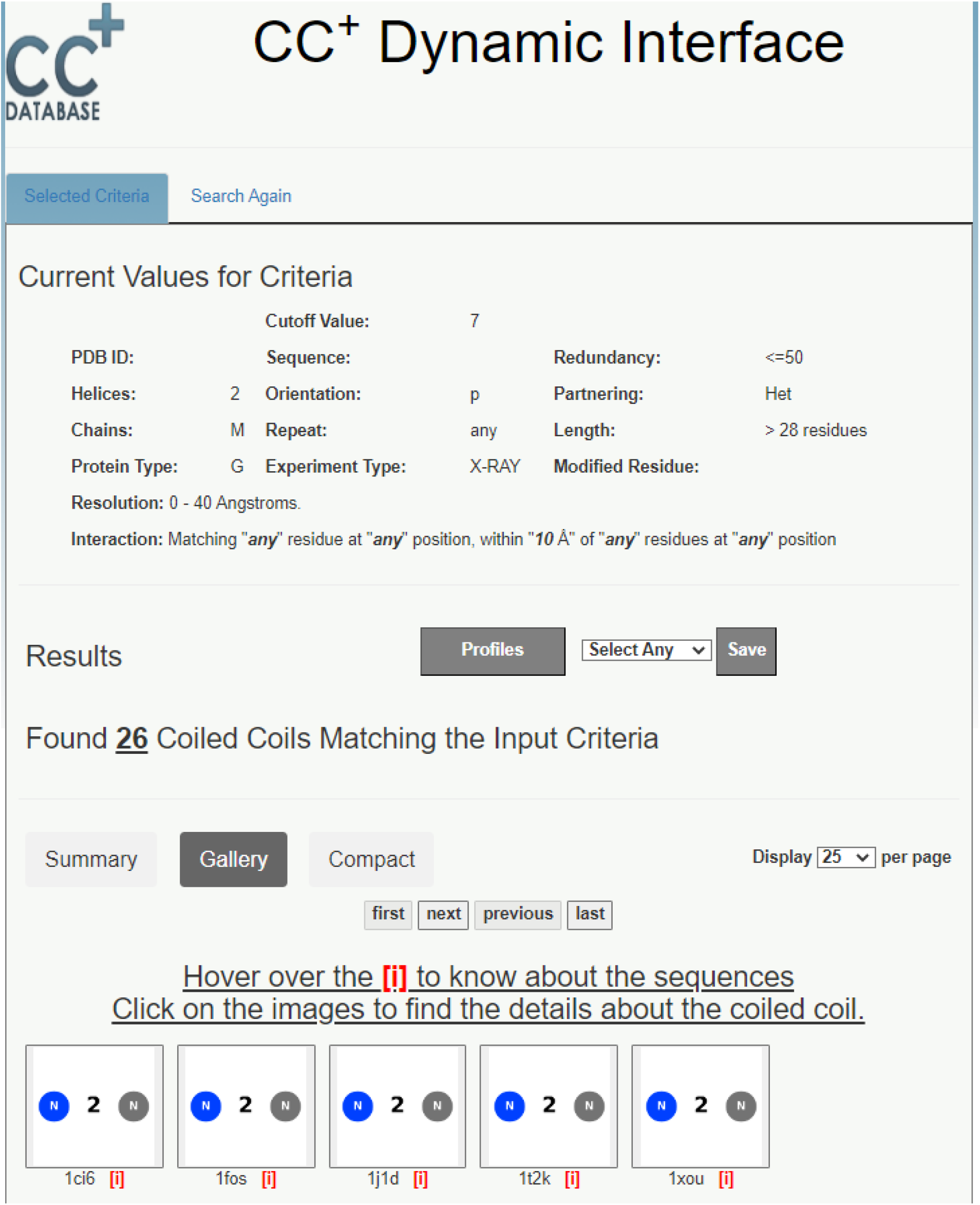
A typical CC^+^ Results Page. In this case, part of the Gallery view of the results is shown at the bottom of the image.

In more detail, the ‘Results’ section has two main features: (a) a list of the returned CCs; and (b) options to download associated data from the search. The subtabs enable displaying the list of CCs in ‘Summary’, ‘Gallery’ or ‘Compact’ forms, with the ‘Gallery’ being the default view. It provides clickable images summarising the architecture of each CC. Users have the option to select the number of CCs displayed, with 25 as the default. In these three displays, the images or PDB codes are also clickable links to a Socket2-based GUI for visualizing the CC in context of the whole structure. This is described in more detail below.

Above these display items, users are given options for downloading data from the search. For instance, the ‘Profiles’ button links to position-specific scoring matrices (PSSMs) for the selected CCs; *i*.*e*., 20x7 tables for the occurrence of each amino acid at the 7 positions of the heptad repeats, which can be normalized internally or using amino-acid frequencies from Uniprot/Swissprot.^33^ In addition, the adjacent drop-down menu and ‘Save’ button allow users to download a flat csv files for the ‘Summary and CC Sequences’, or a zipped file giving the ‘3D-Coordinates’ of the CC regions only in PDB file format. For the latter, the CCs are extended by one residue at each terminus of the constituent helices. These summary, sequence, and coordinate files are provided to facilitate further user-specified analyses and applications of the CC datasets.

Finally, if the database does not provide any hit for a set of search parameters, the ‘Results’ section will display the message ‘No Results Satisfying the Input Criteria.’ In such cases, users are encouraged to redefine their search criteria or to visit the ‘Documentation’ tab.

### 3.4 Improvements on the original 2009 webserver

The database and webserver have undergone significant changes as detailed below.

#### New user interface

The user interface has been updated to ease searches and visualization of results. The ‘Dynamic Interface’ allows users to search for CCs in the PDB or the AlphaFold Protein Structure Database. The ‘Documentation’ tab has been updated accordingly.

#### New search parameters

The updated CC^+^ uses Socket2 to provide broader searches, *e*.*g*. CCs that contain glycine or modified residues. As noted above, users can now specify `Protein Type` as listed in the RCSB PDB, and `Resolution`(in the range 0 – 100 Å) for appropriate `Experiment Types`.

#### Sequences with modified residues

The MODRES record of the PDB file format provides information about the modified residue and the corresponding proteogenic amino acid. Socket2 uses this record allowing CCs that contain non-proteinogenic residues to be identified. CC^+^ can be searched for examples via the ‘Specify Structures’ tab by providing the three-letter code for the non-proteinogenic residue. Typing the first letter of the code activates a drop-down box of non-proteinogenic residues for the user to choose from.

As of January 2023, in the redundant set of CCs found at a cut-off value of 7 Å, 2347 CCs were found with modified residues. Of these, modified methionine (MSE) was the most common accounting for ≈90% of the examples. Others included modified phenylalanine and lysine residues. Interestingly, helical α/β-peptide foldamers (PDB IDs: 2oxj, 2oxk, and 3c3g) from the Gellman lab were identified as CCs by Socket2 and found to have a total of 7 non-proteogenic residues in CC helices, the highest number found to date.^34,35^

#### Ability to query the AlphaFold Protein Structure Database

The new CC^+^ contains CCs found in the AlphaFold2 predicted models from 48 proteomes.^22^ Users can search for CCs from individual proteomes or across all 48 proteomes. However, as the predicted models are for single chains and include only proteogenic residues, some search options of the CCPlus-PDB tab are not available.

When searching CCPlus-AlphaFold, users are given the option to specify ranges of the predicted local distance difference test (`pLDDT Score`). This is a per-residue confidence score for the AlphaFold2 prediction scaled 0 – 100. Scores >90 indicate high confidence, while scores <50 indicate low-confidence predictions. When specified, this option will search for all CCs in CC^+^ with average pLDDT value of the input range. This average is calculated for the residues of the participating helices rather over the whole protein structure.

Note: Currently, CCPlus-AlphaFold only contains CCs that are within the same polypeptide chain; *i*.*e*., only intramolecular CCs. This is a necessary consequence of the publicly available AlphaFold2 models being limited to predictions of tertiary structures. As discussed below, we are working to remedy this by including AlphaFold2-based predictions of quaternary structures.

#### Downloadable resources

As introduced above, users have the option to download the results of CC^+^ as a flat csv file. Also, using Biopython,^28^ PDB files containing only the selected CC regions can be downloaded; in this case the helices are extended by a single residue at each terminus. By clicking on the ‘Profiles’ button of the ‘Results’ page, users can download the raw counts for the occurrence of the 20 standard residues at each heptad position and the Swissprot or internally normalised propensity tables (PSSMs).

#### Visualization of the results

On the ‘Results’ page, each search is presented as ‘Summary’, ‘Gallery’, and ‘Compact’ views. Here, the figure and the PDB ID act as a clickable links to an interactive GUI running Socket2^20^ to visualize and download images of that CC. As detailed in the Socket2 publication,^20^ the GUI gives multiple options for visualizing the component CCs of a structure/model and their associated sequences. Via subtabs, it also provides some analysis of the structures such as CC ‘Register’, ‘Angle Between Helices’, and ‘Core-packing Angles’ for the KIH packing.

#### Regular updates

The webserver is designed to automatically update on the first Thursday of each month. However, updates for the CCPlus-AlphaFold part of the website require manual intervention. We aim to perform these updates regularly to keep the database current with any changes in the AlphaFold Protein Structure Database.^22^

## 4. RESULTS

As described above, the new CC^+^ Database allows searches of CC structures and predicted models from the RSCB PDB and AlphaFold Protein Structure Database, respectively. The new features that we have introduced allow users to search for CCs that are broadly defined or highly specified in terms of structure, sequence, protein type, experimental methods for structure determination, organism, and so on. We have implemented many of these features to give flexibility to users, and because we could not possibly anticipate all searches that users might choose to run. Therefore, here we do not give an exhaustive list or overview of what searches can be done and the data that might be retrieved. Instead, we show a few examples of the shape of the data that CC^+^ contains and how this can be accessed by users.

### 4.1 Distribution of different CCs across dataset

As summarised in the ‘Statistics’ tab and shown in Figure 2, 2-helix CCs make up almost 80% of the total CCs from the PDB, and 87% in the AlphaFold Protein Structure Database. 3-Helix CCs constitute 14% and 11% of the total in PDB and AlphaFold2 datasets, respectively. Whereas 4-helix CCs represent 4% and 1%, respectively. For the CCs from the PDB, intramolecular antiparallel interactions dominate the CCs, although the proportions of parallel and antiparallel interactions even out with increasing numbers of helices in the assemblies. Indeed, the higher-order CC structures in the PDB with >5 helices are mostly parallel, intermolecular assemblies. Because of the nature of the current models in AlphaFold Protein Structure Database, *i*.*e*. they are for tertiary structures only, the predicted CCs are within the same chain and predominantly have antiparallel helices. Although, again in the higher-order CC predictions, at least those captured by the most-stringent Socket2 cut-off of 7 Å, there are 6 parallel and 7 antiparallel predicted 5-helix CCs.

Interestingly, there are some very large CC assemblies with >6 helices, which Socket2 can now identify.^20^ Using the default parameters and ≤70% sequence identity, a total of 24 such CCs were found in the CCPlus-PDB Database. Of these, the true largest assembly has 15 parallel helices, which is in the biological unit of the rotor ring (c subunit) of the proton-dependent ATP synthase (PDB ID, 2wie).^36^ Here, the KIH interactions extend for just eight residues along each helix. This class of protein makes an interesting case study. For instance, the ATP synthase from Ba*cillus pseudofirmus* OF4 has a rotor ring with 13 parallel helices and a slightly longer 11-residue stretch of KIH interactions (PDB ID, 4cbj).^37^ There is a note of caution for Socket2 users too. The program is generally accurate in identifying CCs. However, it can misassign oligomeric state/number of helices. For instance, for the same part from the *E. coli* ATP synthase (PDB ID, 5t4o)^38^ Socket2 indicates a 20-helix CC. However, upon inspection, the structure comprises two concentric 10-helix rings. Similarly, the rotor ring of the mycobacterial ATP synthase (PDB ID, 4v1g)^39^ is a nonomer, but Socket2 identifies it as an 18-mer. The source of the misclassification is that the rings have KIH packing within them and between them; effectively, there are rings of 3-helix bundles, which Socket2 interprets as a single, contiguous, larger ring. This would be difficult to correct for such a small, though interesting, class of structures. Therefore, at this stage, we advise using the visualiser in Socket2 or an external viewer to inspect and verify unusual CC classification from Socket2 and CC^+^.

### 4.2 Comparison of coiled coils in PDB structures and predicted AlphaFold2 models

Using search parameters of 7 Å Socket2 cut-off and ≤70% sequence identity, CCPlus-PDB and CCPlus-AlphaFold currently house ≈12,000 structures and ≈120,000 models, respectively. Such numbers allow good comparison of experimental structures and predicted models to be made; albeit with the caveat that CCPlus-PDB contains both intra- and inter-molecular CCs, whereas, at present, CCPlus-AlphaFold necessarily only has the former.

First, both databases predominantly have CCs (>90%) with canonical (heptad-based) sequence repeats. However, as CC^+^ allows searches of CCs with non-canonical repeats, we asked if the proportions of these differed between the two datasets. Out of 8836 and 90747 CCs, we found 755 and 6116 non-canonical CCs in CCPlus-PDB and CCPlus-AlphaFold datasets (Figure 2), respectively, mirroring the 1:10 size ratio of both datasets.

Next, we compared the experimental and modelled CCs as a function of minimum length binned into five categories: (i) <11, (ii) 11 – 14, (iii) 14 – 21, (iv) 21 – 28, and (v) >28 residues, (**Figure 7**). For the 2-helix structures, the length distributions of the CCs from the two datasets were similar. However, above this, the predicted CCs from the AlphaFold2 dataset tended to be shorter than the corresponding classes from the PDB dataset; this is shown in Figure7B (pink color). We suspect that this is related to the CCs predicted by AlphaFold2 being intra-molecular. Therefore, it will be interesting to see if and how these distributions change when predicted quaternary structures become available.

**FIGURE 7:**
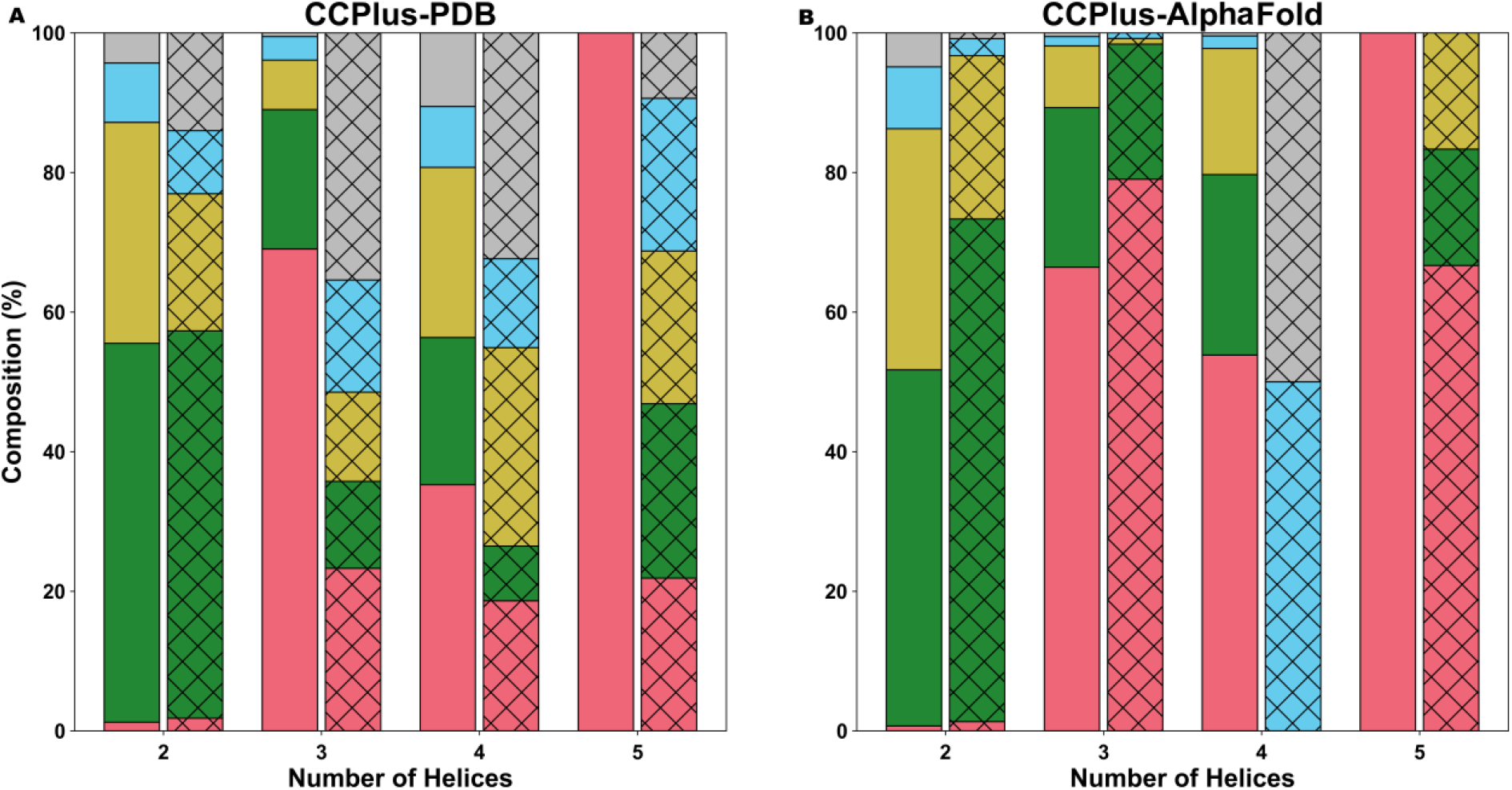
Comparison of CC-length distributions for different CCs in the PDB- and AlphaFold2-derived datasets. Key: solid bars, antiparallel CCs; hatched bars, parallel CCs; pink, CC length <11 residues; green, 11 -14 residues; yellow, 14 – 21 residues; sky-blue, 21 – 28 residues; gray, > 28 residues. CC^+^search parameters: default values, except `Redundancy`: ≤70% sequence identity and `Cutoff`: 7 Å Socket2 cutoff.

Both datasets have examples of CCs with helices >100 residues. For instance, in the PDB dataset, a dimeric CC in a cryo-EM structure of the motor-protein dynein tail-dynactin-BICD2N complex (PDB ID, 5afu)^40^ has a helix spanning 165 residues. And in the AlphaFold2 dataset, a tropomyosin-like protein (Uniprot ID: A0A077ZIM1) from *Trichuris trichiura* is predicted to have a 2-helix antiparallel CC with a helix of 176 residues, although the average pLDDT score is in the confidence range of 70 – 90.

Finally, and to make as direct a comparison as possible, we selected all 2-helix intra-molecular CCs from both datasets. This gave totals of 343 parallel and 3075 antiparallel structures from the CCPlus-PDB, and 5648 and 45514 in the CCPlus-AlphaFold set. Though there is a large difference between the numbers of 2-helix CCs in both datasets, the overall numbers allow for comparative analyses (Supplemental Tables S1-4). In terms of overall amino-acid composition, the sets are very similar (Figures 8A&B). However, the changes in proportions of residues and residue classes at the ***a – g*** sites of the identified heptad repeats of the CCs showed significant differences, as assessed by a z-test (Figures 8C&D). For instance, the proportion of charged residues, Glu and Lys, is lower in CCPlus-AlphaFold dataset at different heptad positions, and compensated by increases in the hydrophobic residues, Ile, Leu, and Phe. This suggests that CCs of the predicted models are less solvent exposed, and/or that many of the predicted structures have the potential to form oligomers. Again, this will have to await analysis of quaternary structure predictions and experimental validations of the AlphaFold2 models.

**FIGURE 8:**
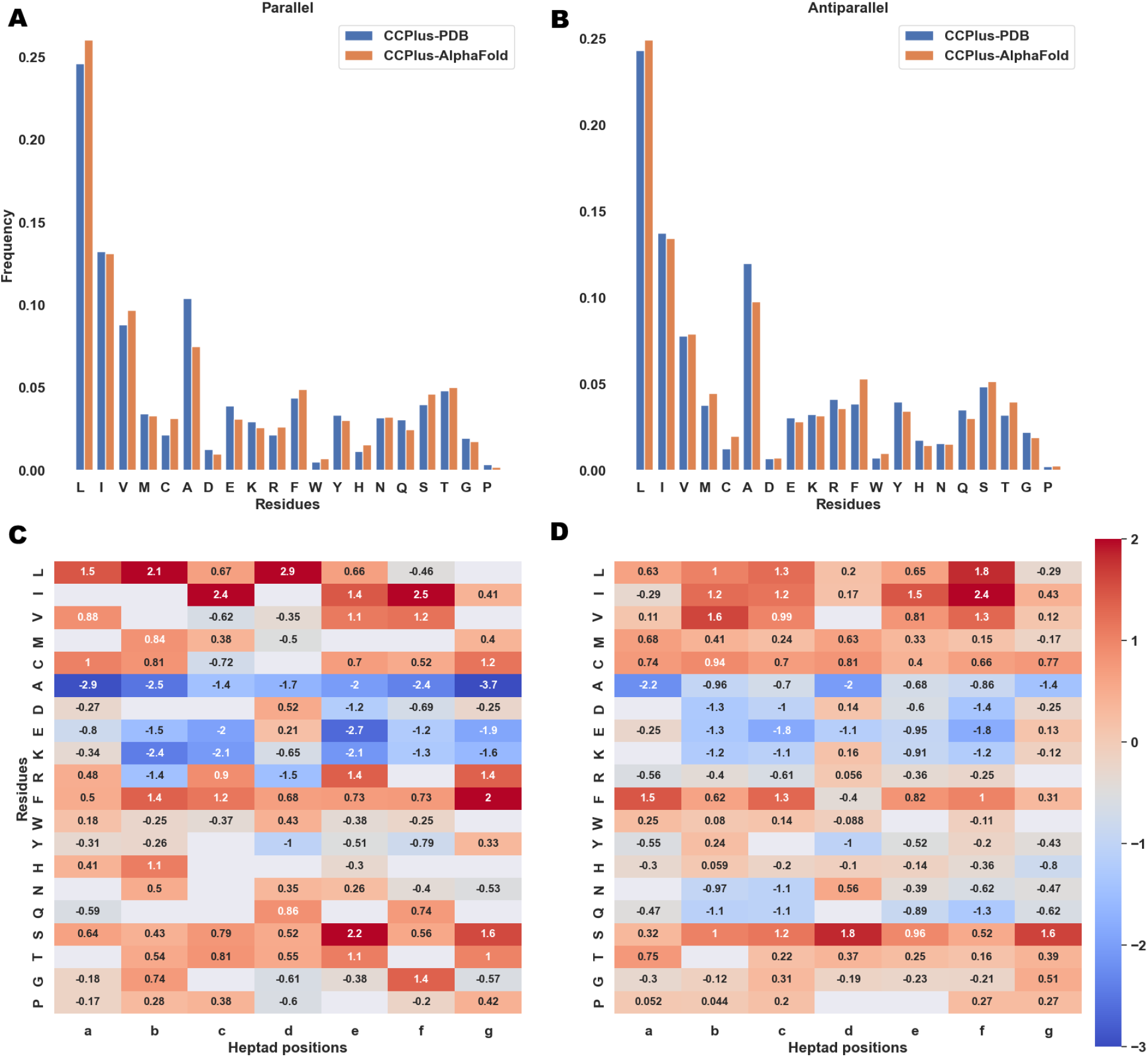
Distribution of 20-proteogenic residues in 2-helix CCs in CCPlus-PDB and CCPlus-AlphaFold datasets. Comparison of frequency distribution of different amino acids at various positions of heptad repeat in (A) parallel and (B) antiparallel 2-helix CCs. Heatmaps for the change of proportion of different residues at heptad positions for (C) parallel and (D) antiparallel 2-helix CCs. The higher the value, the more preferred the residue is in CCPlus-Alphafold dataset. The null hypothesis is that the two datasets have same composition of residues at different position of heptad. Standard amino acids are represented by their one-letter codes. Parallel and antiparallel 2-helix CCs from both datasets were obtained at `Redundancy`: 70% sequence identity and `cutoff`: 7 Å cut-off value. The remaining parameters were set to their default values.

## 5. CONCLUSION

We have described the structure, main features, and a small number of many possible uses of an updated database of coiled-coil (CC) structures and predicted models, the CC^+^ Database. The CCs are found using the program Socket2,^20^ which identifies the signature knobs-into-holes packing between neighboring α helices of CCs. Therefore, it does not rely on, and is not biased by sequence-based CC predictions, which are not always consistent or reliable.^41^ The new CC^+^ Database includes both experimentally derived CCs from the RCSB PDB^14^ and predicted models from 48 genomes using AlphaFold2.^22^ This represents a significant expansion of CC^+^ since its inception in 2009. These two parts of the CC^+^ Database are not directly comparable because, currently, AlphaFold2 predictions are only available for protomers, *i*.*e*., single-chain, tertiary structures. Recent work has predicted homo-oligomeric structure models based on AlphaFold2 for four proteomes.^42^ Interestingly, application of Socket2 to these models indicates that CCs are major enablers of protein-protein interactions, particularly in eukaryotes. However, because that study is distinct from the update of the CC^+^ Database presented here, it will be presented elsewhere.^42^ Nonetheless, the two databases will be linked in the future.

Returning to CC^+^: it is available at http://coiledcoils.chm.bris.ac.uk/CCPlus/Home.html. It can be searched in a wide variety of ways at the sequence and structural levels to generate user-defined datasets. In turn, the identified CCs can be visualized and analyzed in a user-friendly GUI as part of Socket2. There are also options for analyzing datasets as a whole within CC^+^. Alternatively, the datasets (sequences, coordinates, and metadata) can be downloaded in bulk for analysis off-line. Thus, we trust that CC^+^ will be useful to many users interested in an array of CC chemistry, structure, and biology. For instance, from gathering examples of related CC structures for basic biological research to garnering sequence-to-structure/function relationships for underpinning protein design and engineering projects.

## ACKNOWLEDGMENTS

P.K. was supported by the Biotechnology and Biological Sciences Research Council (BBSRC) grant to D.N.W. [BB/R00661X/1]. R.P. was supported by a BBSRC-funded PhD studentship (SWBio DTP). W.M.D. and D.N.W. were funded by a European Research Council Advanced Grant (340764) and a subsequent European Research Council Proof of Concept Grant (787173). D.N.W. was also supported by the BrisSynBio, a BBSRC/Engineering and Physical Sciences Research Council-funded Synthetic Biology Research Centre [BB/L01386X/1]. E.D.L. and H.S. acknowledge support from the European Research Council (ERC) under the European Union’s Horizon 2020 research and innovation program (grant agreement No. 819318), by the Israel Science Foundation (grant no. 1452/18), and by the Abisch-Frenkel Foundation.

## Author contributions

Project conceptualization, PK and DNW. Database and front-end development, PK and DNW. Data analyses: All. Manuscript writing, PK, RP, EDL, and DNW. All authors have read and commented on the manuscript, including the final version.

