## Supplemental Tables 1-4 for "CC^+^: A Searchable Database of Validated Coiled coils in PDB Structures and AlphaFold2 Models"

| | a | b | c | d | e | f | g | Sum |
|---|---|---|---|---|---|---|---|---|
| **L** | 401 | 123 | 127 | 460 | 203 | 177 | 240 | 1731 |
| **I** | 216 | 80 | 58 | 137 | 90 | 70 | 112 | 763 |
| **V** | 144 | 79 | 93 | 123 | 82 | 86 | 109 | 716 |
| **M** | 56 | 18 | 24 | 80 | 38 | 33 | 40 | 289 |
| **C** | 35 | 15 | 33 | 38 | 25 | 25 | 12 | 183 |
| **A** | 170 | 127 | 120 | 154 | 154 | 172 | 179 | 1076 |
| **D** | 21 | 48 | 41 | 13 | 74 | 74 | 49 | 320 |
| **E** | 64 | 100 | 95 | 47 | 138 | 123 | 129 | 696 |
| **K** | 48 | 91 | 83 | 54 | 100 | 95 | 96 | 567 |
| **R** | 35 | 76 | 56 | 62 | 56 | 78 | 72 | 435 |
| **F** | 72 | 35 | 41 | 61 | 54 | 54 | 34 | 351 |
| **W** | 9 | 11 | 21 | 16 | 30 | 17 | 16 | 120 |
| **Y** | 55 | 30 | 35 | 61 | 55 | 49 | 41 | 326 |
| **H** | 19 | 14 | 20 | 21 | 31 | 25 | 21 | 151 |
| **N** | 52 | 37 | 36 | 30 | 40 | 63 | 43 | 301 |
| **Q** | 50 | 59 | 52 | 31 | 70 | 54 | 82 | 398 |
| **S** | 65 | 71 | 73 | 71 | 71 | 91 | 60 | 502 |
| **T** | 79 | 45 | 49 | 61 | 55 | 72 | 51 | 412 |
| **G** | 32 | 46 | 49 | 49 | 56 | 59 | 41 | 332 |
| **P** | 6 | 7 | 7 | 30 | 13 | 14 | 2 | 79 |

**Supplemental Table S1**: 20 x 7 matrix of the occurrence of the 20 proteinogenic residues at each position of the heptad repeat for 343 parallel 2-helilx CCs in CCPlus-PDB. The last column lists the total number of occurrences of the residue in the dataset. CCs were obtained at Redundancy of 70% sequence identity or less, and with a Socket2 cut-off of 7 Å. The remaining parameters were set to their default values.

| | a | b | c | d | e | f | g | Sum |
|---|---|---|---|---|---|---|---|---|
| **L** | 4066 | 1148 | 1267 | 4656 | 1840 | 1161 | 2102 | 16240 |
| **I** | 2299 | 739 | 765 | 1419 | 766 | 651 | 954 | 7593 |
| **V** | 1304 | 709 | 818 | 1161 | 810 | 758 | 969 | 6529 |
| **M** | 633 | 219 | 275 | 511 | 336 | 297 | 490 | 2761 |
| **C** | 213 | 97 | 131 | 435 | 193 | 141 | 190 | 1400 |
| **A** | 2006 | 1410 | 1259 | 2068 | 1254 | 1512 | 1784 | 11293 |
| **D** | 119 | 798 | 672 | 199 | 621 | 854 | 484 | 3747 |
| **E** | 513 | 1230 | 1274 | 580 | 1393 | 1452 | 1200 | 7642 |
| **K** | 542 | 1000 | 961 | 319 | 1064 | 1239 | 870 | 5995 |
| **R** | 693 | 783 | 788 | 243 | 915 | 895 | 766 | 5083 |
| **F** | 645 | 452 | 519 | 1070 | 525 | 507 | 526 | 4244 |
| **W** | 124 | 103 | 149 | 234 | 159 | 166 | 137 | 1072 |
| **Y** | 667 | 276 | 409 | 887 | 466 | 396 | 435 | 3536 |
| **H** | 296 | 227 | 280 | 318 | 275 | 303 | 348 | 2047 |
| **N** | 263 | 684 | 587 | 364 | 574 | 705 | 474 | 3651 |
| **Q** | 587 | 833 | 807 | 378 | 993 | 937 | 936 | 5471 |
| **S** | 810 | 777 | 722 | 736 | 844 | 938 | 638 | 5465 |
| **T** | 538 | 613 | 611 | 738 | 728 | 729 | 589 | 4546 |
| **G** | 373 | 641 | 495 | 436 | 682 | 734 | 490 | 3851 |
| **P** | 39 | 123 | 70 | 106 | 84 | 116 | 27 | 565 |

**Supplemental Table S2**: 20 x 7 matrix of the occurrence of the 20 proteinogenic residues at each position of the heptad repeat for 3075 antiparallel 2-helilx CCs in CCPlus-PDB. The last column lists the total number of occurrences of the residue in the dataset. CCs were obtained at Redundancy of 70% sequence identity or less, and with a Socket2 cut-off of 7 Å. The remaining parameters were set to their default values.

| | a | b | c | d | e | f | g | Sum |
|---|---|---|---|---|---|---|---|---|
| **L** | 7354 | 2722 | 2496 | 9043 | 3667 | 2931 | 4117 | 32330 |
| **I** | 3706 | 1462 | 1574 | 2367 | 1909 | 1823 | 2027 | 14868 |
| **V** | 2741 | 1523 | 1598 | 2093 | 1693 | 1772 | 1891 | 13311 |
| **M** | 937 | 508 | 525 | 1284 | 670 | 542 | 785 | 5251 |
| **C** | 898 | 447 | 464 | 685 | 604 | 559 | 494 | 4151 |
| **A** | 2114 | 1838 | 1931 | 2266 | 2154 | 2375 | 2165 | 14843 |
| **D** | 287 | 858 | 784 | 379 | 982 | 1103 | 780 | 5173 |
| **E** | 882 | 1544 | 1359 | 899 | 1722 | 1811 | 1752 | 9969 |
| **K** | 736 | 1189 | 1117 | 778 | 1202 | 1317 | 1257 | 7596 |
| **R** | 741 | 1113 | 1226 | 684 | 1310 | 1360 | 1572 | 8006 |
| **F** | 1388 | 931 | 1000 | 1281 | 1112 | 1109 | 1083 | 7904 |
| **W** | 207 | 153 | 314 | 409 | 424 | 232 | 245 | 1984 |
| **Y** | 864 | 503 | 610 | 794 | 822 | 649 | 786 | 5028 |
| **H** | 446 | 480 | 388 | 400 | 460 | 445 | 396 | 3015 |
| **N** | 917 | 791 | 676 | 635 | 755 | 985 | 610 | 5369 |
| **Q** | 700 | 1052 | 941 | 799 | 1215 | 1112 | 1360 | 7179 |
| **S** | 1307 | 1408 | 1518 | 1415 | 1777 | 1703 | 1422 | 10550 |
| **T** | 1426 | 949 | 1078 | 1244 | 1217 | 1245 | 1128 | 8287 |
| **G** | 502 | 1008 | 859 | 701 | 873 | 1351 | 564 | 5858 |
| **P** | 56 | 189 | 209 | 365 | 205 | 191 | 138 | 1353 |

**Supplemental Table S3**: 20 x 7 matrix of the occurrence of the 20 proteinogenic residues at each position of the heptad repeat for 5648 parallel 2-helilx CCs in CCPlus-AlphaFold. The last column lists the total number of occurrences of the residue in the dataset. CCs were obtained at Redundancy of 70% sequence identity or less, and with a Socket2 cut-off of 7 Å. The remaining parameters were set to their default values.

| | a | b | c | d | e | f | g | Sum |
|---|---|---|---|---|---|---|---|---|
| **L** | 65403 | 20591 | 22970 | 73413 | 30119 | 22134 | 32105 | 266735 |
| **I** | 35277 | 14463 | 14706 | 22671 | 15225 | 15466 | 15826 | 133634 |
| **V** | 20735 | 14754 | 15205 | 18357 | 14433 | 14757 | 15362 | 113603 |
| **M** | 11714 | 4370 | 4914 | 9672 | 5985 | 4954 | 7252 | 48861 |
| **C** | 5273 | 3496 | 3546 | 8939 | 3918 | 3692 | 4682 | 33546 |
| **A** | 25639 | 20696 | 18786 | 27008 | 17980 | 21588 | 24573 | 156270 |
| **D** | 1909 | 10173 | 8653 | 3487 | 8304 | 10191 | 6979 | 49696 |
| **E** | 7392 | 17062 | 16699 | 6049 | 19550 | 18544 | 18994 | 104290 |
| **K** | 8328 | 13546 | 13185 | 5422 | 14507 | 16559 | 13283 | 84830 |
| **R** | 9397 | 11766 | 11404 | 3951 | 13435 | 13353 | 11997 | 75303 |
| **F** | 13936 | 8549 | 11050 | 15706 | 10020 | 10193 | 8899 | 78353 |
| **W** | 2596 | 1821 | 2692 | 3431 | 2524 | 2337 | 2088 | 17489 |
| **Y** | 9009 | 4935 | 6425 | 11129 | 6083 | 5701 | 5808 | 49090 |
| **H** | 3861 | 3772 | 4082 | 4710 | 3962 | 3905 | 3621 | 27913 |
| **N** | 4075 | 9003 | 7253 | 7172 | 8061 | 9577 | 6322 | 51463 |
| **Q** | 7974 | 11147 | 10792 | 5850 | 13448 | 11687 | 13197 | 74095 |
| **S** | 13540 | 14628 | 13992 | 16338 | 15307 | 15758 | 13633 | 103196 |
| **T** | 10409 | 9927 | 10270 | 12524 | 11908 | 11692 | 10055 | 76785 |
| **G** | 5052 | 10070 | 8591 | 6333 | 10093 | 10952 | 8776 | 59867 |
| **P** | 749 | 2069 | 1543 | 1734 | 1245 | 2406 | 1020 | 10766 |

**Supplemental Table S4**: 20 x 7 matrix of the occurrence of the 20 proteinogenic residues at each position of the heptad repeat for 45514 antiparallel 2-helilx CCs in CCPlus-AlphaFold. The last column lists the total number of occurrences of the residue in the dataset. CCs were obtained at Redundancy of 70% sequence identity or less, and with a Socket2 cut-off of 7 Å. The remaining parameters were set to their default values.
